# Early life Oxytocin treatment Attenuates Seizure Susceptibility in Male, but not Female, *Fmr1*-KO Mice

**DOI:** 10.64898/2026.06.17.732971

**Authors:** Jasmine Chavez, Julie C. Lauterborn, Gary Lynch, Christine M. Gall

**Author notes:** **Corresponding author:** Christine M. Gall, Gillespie Neuroscience Research Facility, 837 Health Science Road, Rm 3226, University of California, Irvine, CA 92697. **Author emails:** Jasmine Chavez, Julie Lauterborn, Gary Lynch, Christine M. Gall.

## Abstract

Fragile X syndrome (FXS) is the leading inherited cause of intellectual disability, and is frequently accompanied by seizures. Early-life treatment with the hormone oxytocin (OXT) improves social behavior and cognitive function in rodent models of autism with intellectual disability, including FXS, but potential OXT treatment effects on seizure susceptibility have not been evaluated. Here we tested, in both sexes, if intranasal OXT (iOXT) or saline (iSAL) during the second postnatal week reduces audiogenic seizures (AGS) in the *Fmr1*-Knockout (KO) mouse model of FXS. OXT given daily from postnatal day (P) 7 to P13 significantly reduced the incidence and severity of AGS and the latency to seize in adult male *Fmr1*-KOs. Female KOs exhibited less severe seizures that were unaffected by treatment. Wild type mice did not exhibit AGS independent of treatment. To test if antiepileptic effects of iOXT are age-dependent, a separate cohort received iOXT daily from P30 to P36. Male KOs receiving later treatments exhibited robust seizures that were comparable between OXT- and SAL-treatment groups, suggesting that OXT’s enduring antiepileptic effects are confined to early postnatal treatments. Tests of acute OXT effects in adulthood demonstrated an attenuation of male *Fmr1*-KO AGS at testing 30-60 min and 1 day post-treatment but these effects were not evident 15 days later. These findings reveal marked sex differences in the propensity for audiogenic seizures in *Fmr1*-KO mice and demonstrate that early-life OXT treatment mitigates seizure susceptibility in males FXS model mice.

## Introduction

Fragile X syndrome (FXS) is the most prevalent inherited cause of intellectual disability, with relatively high comorbidity for autism spectrum disorder (ASD) (Kaufmann et al., 2017). FXS arises from a CGG triplet repeat expansion in the *Fmr1* gene on the X chromosome, resulting in the silencing of fragile X messenger ribonucleoprotein 1 (FMRP) expression (Pieretti et al., 1991; Verkerk et al., 1991). As a result, learning impairments, sensory hyperexcitability, anxiety, and sociability deficits are known to occur (Hagerman and Hagerman, 2002; Lozano et al., 2014). Symptoms also include a high predisposition for seizures (Berry-Kravis, 2002; Incorpora et al., 2002). Approximately 10–20% of individuals with FXS experience seizures, with a greater incidence (31%) in those cases exhibiting ASD (Garcia-Nonell et al., 2008; Qiu et al., 2008; Berry-Kravis et al., 2010). Although often outgrown after childhood, seizure activity in FXS can leave enduring disruptions in neuronal excitability and developmental trajectories with significant consequences for behavior (Hodges et al., 2019; Berry-Kravis et al., 2021). Thus, the need for effective therapeutics is vital.

Oxytocin (OXT), a hypothalamic neuropeptide, plays a pivotal role in regulating social behavior and parturition (Ferguson et al., 2000; Winslow and Insel, 2002) but has also emerged as a potential therapeutic for synaptic and behavioral impairments in neurodevelopmental disorders such as ASD and FXS (Hall et al., 2012; Peñagarikano et al., 2015; Pan et al., 2022; Chavez et al., 2026). Both clinical and experimental models of autism and FXS demonstrate reduced OXT levels (Modahl et al., 1998; Francis et al., 2014), and OXT-deficient mice exhibit autistic-like behaviors along with increased seizure susceptibility (Sala et al., 2011; Leonzino et al., 2016). These findings have prompted investigation into the anticonvulsant properties of OXT, with accumulating evidence demonstrating its capacity to attenuate seizure activity in mouse models of ASD (Sala et al., 2011; Tyzio et al., 2014; Leonzino et al., 2016) and temporal lobe epilepsy (Erbas et al., 2013; Sahin et al., 2022; Chen et al., 2023).

Prior work suggests that heightened seizure susceptibility in FXS reflects an imbalance between excitatory-inhibitory activity due to impaired GABAergic synaptic function during postnatal development, as has been observed in the mouse model (Rubenstein and Merzenich, 2003; Belmonte and Bourgeron, 2006; Adusei et al., 2010; Paluszkiewicz et al., 2011; He et al., 2014). OXT has been shown to directly influence the maturation of inhibitory systems by promoting the timely upregulation of the potassium and chloride cotransporter (KCC2), which lowers intracellular chloride levels and enables GABA_A_ receptor-mediated inhibition (Tyzio et al., 2006; Tyzio et al., 2014; Leonzino et al., 2016). These findings implicate OXT deficiencies in the pathophysiology of FXS and suggest that OXT treatments may have therapeutic benefits in mitigating seizure activity.

The present studies tested if early postnatal OXT treatment attenuates seizures in the *Fmr1*-Knockout (KO) mouse model of FXS, which exhibits the main behavioral and physiological features of this condition as well as heightened audiogenic seizure (AGS) susceptibility (Musumeci et al., 2000; Chen and Toth, 2001; Kazdoba et al., 2014). A predisposition for AGS is one of the most robust and well-characterized phenotypes in *Fmr1*-KOs: Seizures can be induced by brief acoustic stimulation (120 dB) and this response has been extensively studied to probe neural circuits and molecular mechanisms underlying epilepsy in FXS (Musumeci et al., 2000; Chen and Toth, 2001; Yan et al., 2004). Given that FXS influences both males and females (Hunter et al., 2014), effects of early OXT treatment on AGS were evaluated in both sexes.

Our results show that in *Fmr1*-KO mice, OXT treatment during the second postnatal week, but not during adolescence, attenuated AGS in adult males whereas the milder seizure behavior exhibited by females was not altered by OXT treatment. These results demonstrate that early-life OXT treatment can lead to an enduring, and potentially life-long, elimination of seizure susceptibility in male *Fmr1*-KOs, and potentially individuals with FXS. The absence of OXT treatment effects in females raises the possibility that OXT actions, and potentially neuronal OXT signaling, is sexually dimorphic.

## METHODS

### Animals

Male and female wild-type (WT) and *Fmr1*-KO mice (on the sighted FVB129 background) reared in-house were used. After weaning at P21, mice were group-housed with same-sex littermates (3-5 per cage). Mice were maintained in rooms at 68°F and 55% humidity with 12 hr on/12 hr off light cycle (lights on 6AM) and food and water provided *ad libitum*. Behavioral testing was conducted from 10 AM to 4 PM. Protocols were approved by the Institutional Animal Care and Use Committee at the University of California, Irvine and were consistent with the NIH Guide for the Care and Use of Laboratory Animals.

### Oxytocin Treatments (Peñagarikano et al., 2015; Chavez et al., 2026)

WT and *Fmr1*-KO mice of both sexes were given daily intranasal treatments (2 µL per nostril) with OXT (1 ug/uL) or 0.9% NaCl (SAL) using a P10 micropipette Gilson pipette (Pipetman P10, 1-10µL) (**Fig. 1**). Stock solutions of 10mg/mL synthetic human OXT (CellSciences, CRO300A) were diluted with SAL, to obtain the 1ug/uL treatment solution. Prior to treatment, all pups in a litter were briefly relocated to a standard holding cage alongside their home cage and individually returned to the home cage (with dam) after inhalation of the treatment solution was observed. Experimenter’s gloves and holding cage bedding were changed between litters. Handling was consistent across all treatment groups and lasted roughly 30 s per mouse. Groups of mice were treated daily for 7 days from P7 to P13 or from P30 to P36. All pups in a given litter received the same treatment. After treatment of the younger cohort on P13, the pups remained untouched with their littermates and dam until weaning at P21 and thereafter remained group-housed with same-sex littermates until testing.

**Figure 1.**
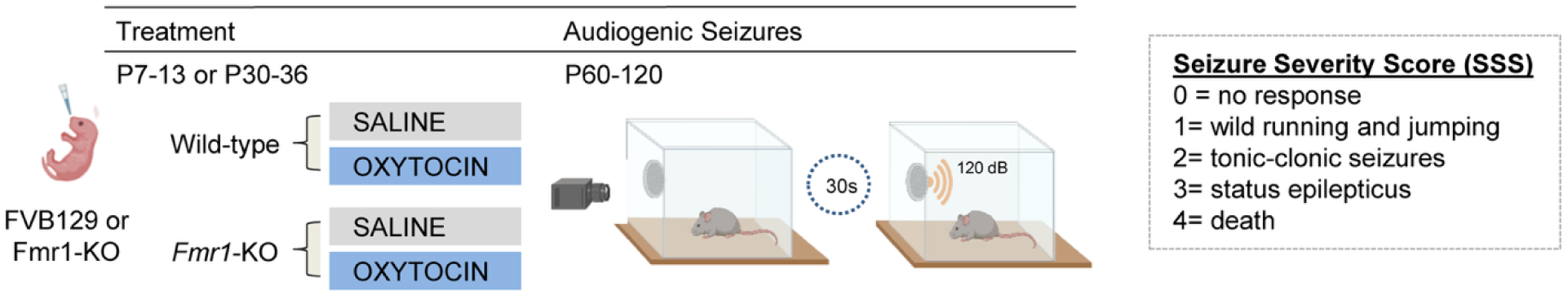
Experimental Paradigm. FVB129-wildtype (WT) and *Fmr1-*KO mice were given daily intranasal oxytocin (OXT) or saline (SAL) treatments from postnatal day (P) 7 to P13 or P30 to P36 and audiogenic seizure assays were performed in adulthood (P60-120). Mice were given 30 s for habituation in the test chamber and then a 120dB auditory stimulus was introduced for 45 s. Seizure responses were classified using the seizure severity score (SSS).

To assess the effects of acute OXT treatment in adults, a separate cohort of 2-3 mo old naive *Fmr1*-KO and WT mice were given OXT (5 mg/kg) or 0.9% NaCl (SAL-vehicle) via intraperitoneal injection (IP); the IP route was used to assure effective administration of the intended amount and because of the short interval between OXT/SAL administration and initial testing. These mice were assessed for AGS 30-60 min after treatment and again 24hrs and 15 days later.

### Audiogenic Seizures (Musumeci et al., 2000; Ding et al., 2014; Gonzalez et al., 2019)

The incidence and severity of AGS were assessed as described (Musumeci et al., 2000; Ding et al., 2014; Gonzalez et al., 2019). Briefly, individual mice were placed inside a plexiglass arena (60 cm x 60 cm floors x 30 cm walls) containing a door alarm (GE 50246 personal security alarm) mounted onto one inside wall; the arena was then covered by a lid. The mouse was given 30 s to explore the arena and then the 120 dB alarm was sounded for 45 s (Guo et al., 2016; Gonzalez et al., 2019). Analysis of the animal’s behavioral response to the alarm was captured using a video camera and scored using the following seizure severity score (SSS): 0 = no response; 1 = wild running and jumping; 2 = tonic-clonic seizures; 3 = status epilepticus; and 4 = death (Musumeci et al., 2007). Sound intensity was calibrated prior to the alarm’s experimental use with a sound level meter (AppRover, 2020; Boulogne-Billancourt, France). Prior to behavioral testing, all adult female mice were evaluated for estrous stage by vaginal lavage (McLean et al., 2012; Cora et al., 2015).

### Statistical analysis

All significance testing was performed using GraphPad Prism 6.0 (San Diego, CA). A nonparametric Kruskal Wallis test with post-hoc Dunn’s analysis was used for genotype and treatment comparisons. Kaplan–Meier survival analysis was used to assess effects on latency to seizure onset, where the event was defined as the post-alarm latency at which the animal entered the first seizure stage on the SSS (wild running and jumping); for this analysis the plots show the proportion of mice remaining seizure-free over time. Statistical significance was assessed at p<0.05. Graphs all show group mean ± standard error mean (SEM) values.

## RESULTS

### Early-age OXT treatment attenuates audiogenic seizures in adult male Fmr1-KO mice

To test if early postnatal OXT treatment influences the predisposition for AGS, WT and *Fmr1*-KO mice of both sexes were given intranasal treatments with OXT (2µL of 1ug/uL per naris) or SAL (iOXT or iSAL, respectively) daily from P7 to P13. In adulthood, the mice were assessed for the expression of AGS with behavior ranked using the SSS (Musumeci et al., 2007) (**Fig. 1 and 2A**).

**Figure 2.**
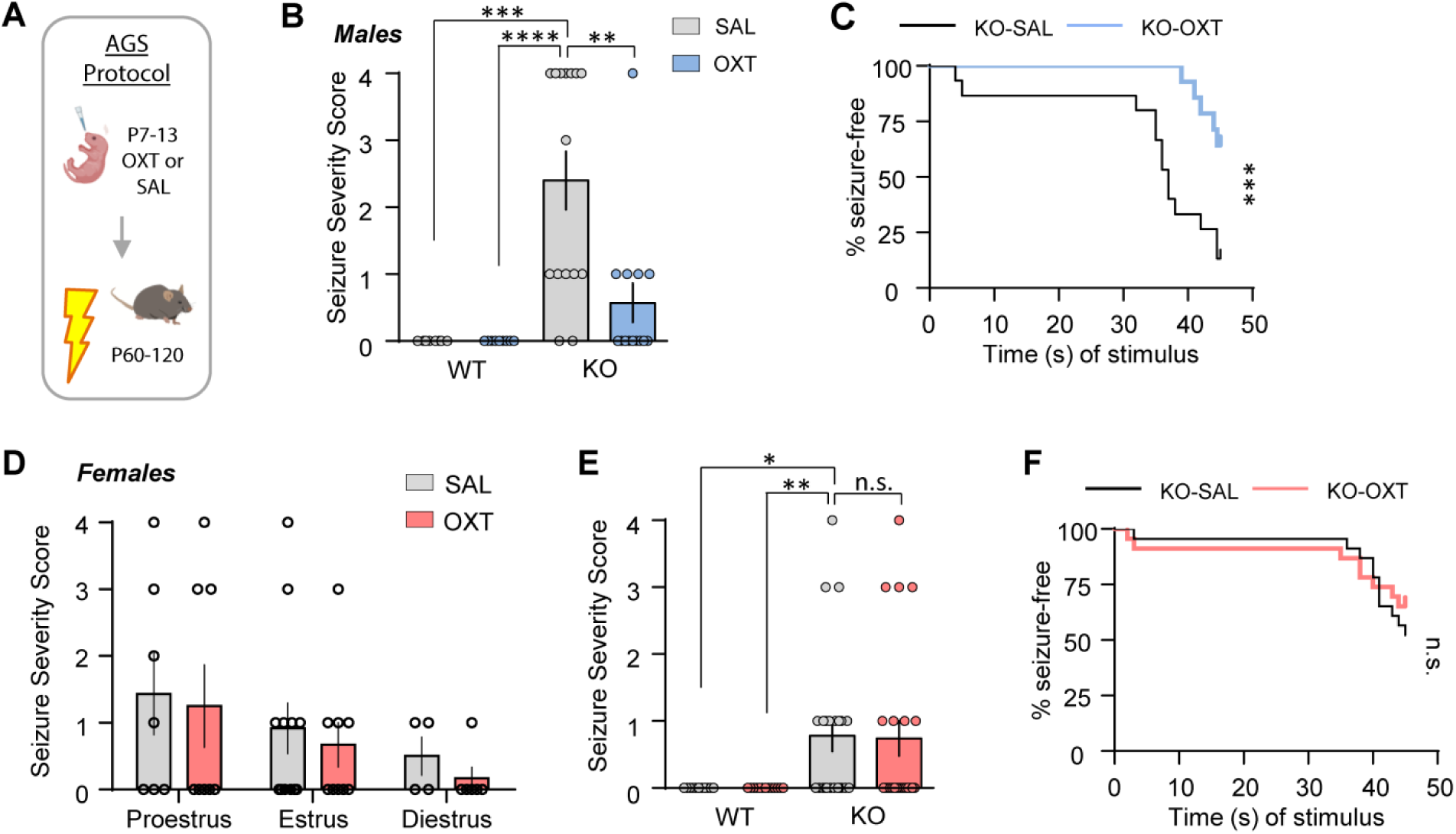
Early-life oxytocin treatment mitigates audiogenic seizures (AGS) in adult male, but not female, *Fmr1-*KO mice. (**A**) Schematic of AGS paradigm for mice treated with oxytocin (OXT) or saline (SAL) from P7-P13 and tested as adults. (**B**) In males, seizure activity was absent in wild-type (WT) mice but robust in the SAL-treated Fmr1-KO (KO) group. Seizure activity was severely reduced or eliminated by OXT treatment (H=25.54, p<0.0001; WT-SAL n=8, WT-OXT n=9, KO-SAL n=15, KO-OXT n=14). (**C**) Latency to seizure onset was significantly greater in *Fmr1*-KOs given OXT vs SAL (Log-rank= 10.88, p=0.001; Hazard ratio (KO-SAL) = 4.56 (95% CI= 2.061 to 14.1; KO-SAL n=13, KO-OXT n=5)). (**D**) In female *Fmr1*-KOs (±OXT) AGS activity was comparable across estrus stages (K-W=3.28, p=0.66). (**E**) Among females, seizure activity was absent in WTs and modest in KOs, but there was no effect of OXT treatment (H=13.73, p=0.003; WT-SAL n=11, WT-OXT n=13, KO-SAL n=23, KO-OXT n=23). (**F**) The latency to seizure onset did not differ between female *Fmr1*-KO groups (Log-rank= 0.46, p=0.50; Hazard ratio (KO-SAL)= 1.36 (95% CI= 0.56 to 3.37; KO-SAL n=11, KO-OXT n=8). Legend in D applies to panels E and F. *Statistics*: Kruskal–Wallis (H) with Dunn’s post-hoc comparisons *(B, D, E)* and Log-rank (Mantel-Cox) test with hazard ratio (logrank) (*C, F)* indicated as: *p<0.05, **p<0.01, ***p<0.001, and ****p<0.0001.

In males, neither WT group displayed AGS (WT-SAL: 0.0 ± 0.0; WT-OXT: 0.0 ± 0.0). In contrast, SAL-treated *Fmr1*-KOs exhibited robust seizure behavior with approximately half of the group going into cardiac arrest. The seizure response was significantly attenuated but not fully eliminated in male KOs that received early iOXT (H=25.54, *p*<0.0001; KO-SAL: 2.40 ± 0.43; KO-OXT: 0.57 ± 0.29, Kruskal-Wallis) (**Fig. 2B**). Moreover, the latency to enter the first seizure stage, defined as wild running or jumping, was significantly longer in OXT-treated *Fmr1*-KOs relative to mutants given iSAL (Log-rank (Mantel-Cox) test=10.88; p=0.001) (**Fig. 2C**).

It is known that in females, seizure susceptibility is influenced by menstrual cycle (Herzog, 2008; Verrotti et al., 2012) and that estrogen and progesterone exhibit proconvulsant and anticonvulsant properties, respectively (Foldvary-Schaefer and Falcone, 2003; Herzog, 2008). Therefore, analyses of AGS in females first tested if estrous stage influenced the severity of seizure behavior and response to OXT treatments. Independent of treatment, female *Fmr1*-KOs in proestrus and estrus stages exhibited somewhat more robust seizure behavior than was the case for those in diestrus (including diestrus and metestrus together) but there were no statistical differences across the stages (H=3.28, p=0.66; Kruskal-Wallis) (**Fig. 2D**). Thus, the results for females at different estrous stages were pooled for further statistical analyses. As shown in Figure 2E, in females, as in males, the WT groups did not display seizure responses (WT-SAL: 0.0 ± 0.0; WT-OXT: 0.0 ± 0.0). However, in contrast to males, female *Fmr1*-KOs exhibited only modest seizure activity (Males vs. Females (KO-SAL): Mann-Whitney U =74.50, p=0.002) and there was no effect of early OXT treatment on the group seizure severity score (KO-SAL: 0.78 ± 0.23; KO-OXT: 0.74 ± 0.26, Kruskal-Wallis; KO-SAL vs KO-OXT: p≥0.99, Dunn’s post hoc) (**Fig. 2E**). Similarly, the latency to the first seizure response did not differ between OXT- and SAL-treated *Fmr1*-KO females (Log-rank (Mantel-Cox) test = 0.46; p=0.46) (**Fig. 2F**).

These results demonstrate clear sex differences in the expression of AGS in full *Fmr1*-KOs and that the anti-epileptogenic effects of early-life OXT treatment are limited to males. Given that early-age iOXT was only found to alleviate seizure responses in males, the following studies used males only.

### Adolescent OXT treatment does not attenuate AGS in male Fmr1-KO mice

Previous work has demonstrated that OXT is involved in orchestrating critical neurodevelopmental events early in life, particularly through influences on the maturational, negative shift in the GABA receptor chloride reversal potential (Sala et al., 2011; Tyzio et al., 2014; Leonzino et al., 2016). Notably, these GABAergic developmental processes are delayed in *Fmr1*-KO mice and this effect is thought to contribute to altered circuit excitability, network dysfunction, and heightened seizure susceptibility in the model (Adusei et al., 2010; Paluszkiewicz et al., 2011; He et al., 2014; Van der Aa and Kooy, 2020). The developmental window during which OXT is known to influence GABAergic transmission closely aligns with the early postnatal period targeted in the present study, suggesting that the efficacy of OXT treatment may be tightly constrained by developmental timing. To evaluate this possibility, male *Fmr1*-KO and WT mice were given intranasal OXT or SAL treatments during adolescence (P30–P36) and then assessed for AGS susceptibility in adulthood (i.e., ≥ P60) (**Fig 3A**). Consistent with our previous findings, WT mice receiving either treatment did not exhibit AGS (WT-SAL: 0.0 ± 0.0; WT-OXT: 0.0 ± 0.0). In marked contrast, *Fmr1*-KOs given OXT in adolescence displayed severe seizures that were comparable to KOs given SAL (KO-SAL vs KO-OXT: p≥0.99, Dunn’s post hoc; KO-SAL: 2.4 ± 0.43; KO-OXT: 0.57 ± 0.29; H=20.55, *p*=0.0001, Kruskal-Wallis) (**Fig. 3B**). Seizure latency was unaffected by treatment in *Fmr1*-KO mice (Log-rank (Mantel-Cox) test = 0.88; p=0.35) (**Fig. 3C**). Together, these findings indicate the presence of a developmentally restricted window during which OXT treatment can produce enduring neuroprotective effects.

**Figure 3.**
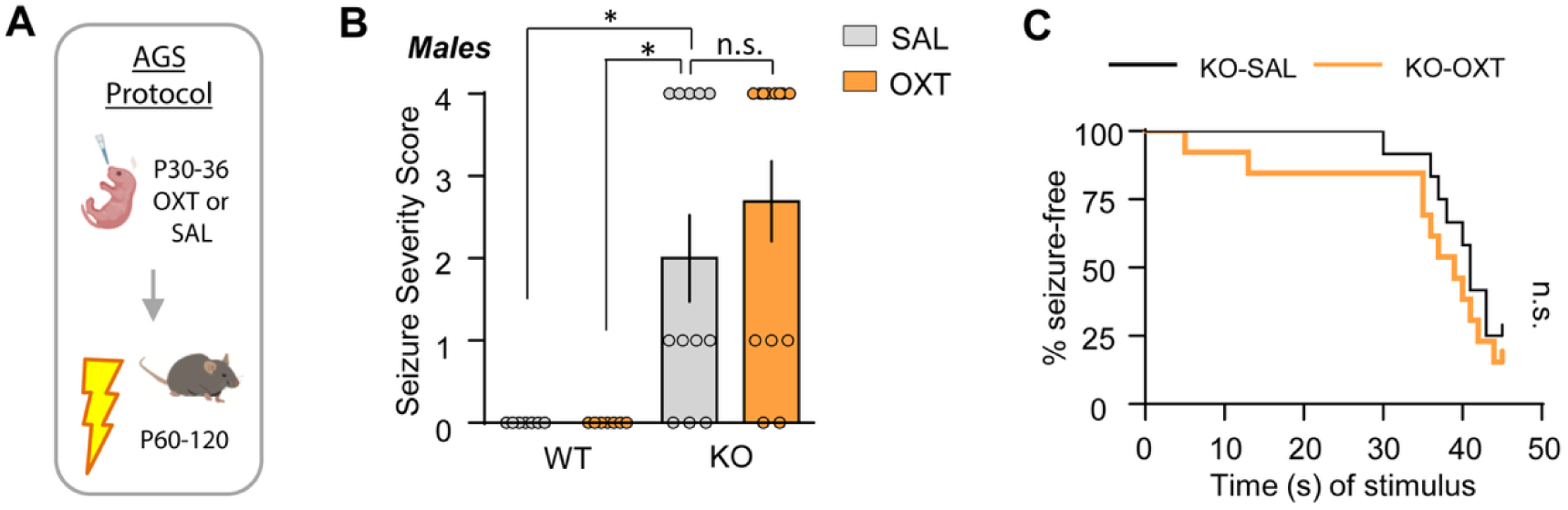
Adolescent oxytocin treatment did not eliminate audiogenic seizure (AGS) responses in male *Fmr1*-KOs. (**A**) Schematic of AGS paradigm: mice were treated daily with oxytocin (OXT) or saline (SAL) from postnatal days (P) 30-36 and tested as adults. (**B**) Male wild-type (WT) mice treated with OXT or SAL did not display AGS, whereas both SAL- and OXT-treated *Fmr1*-KOs exhibited robust AGS (H=20.55, p=0.0001; WT-SAL n=7, WT-OXT n=7, KO-SAL n=12, KO-OXT n=13). (**C**) Latency to seizure onset was comparable between KO-SAL and KO-OXT groups (Log-rank= 0.88, p=0.35; Hazard ratio (KO-SAL)= 0.67 (95% CI= 0.27 to 1.56; KO-SAL n=9, KO-OXT n=11). *Statistics*: Kruskal–Wallis (H) with Dunn’s post-hoc comparisons *(B)* and Log-rank (Mantel-Cox) test with hazard ratio (logrank) (*C)* indicated as: *p<0.05.

### Acute OXT treatment transiently reduces seizure susceptibility in adult male Fmr1-KO mice

Acute OXT treatment reportedly reduces seizure activity in mouse models of neurological disorders, including autism (Sala et al., 2011; Erfanparast et al., 2017; Panaitescu et al., 2018; Chen et al., 2023) but this possibility has not been tested in *Fmr1*-KOs. Therefore, to address this question, naïve adult male *Fmr1*-KO and WT mice were given a single IP injection of OXT (5mg/kg) or SAL 30 to 60 min prior to AGS testing (**Fig. 4A**). WT mice did not exhibit seizures regardless of treatment (WT-SAL: 0.0 ± 0.0; WT-OXT: 0.0 ± 0.0), whereas SAL-treated *Fmr1*-KOs displayed robust seizures. AGS behavior was markedly attenuated in *Fmr1*-KOs given acute OXT treatment (H=14.11, *p*=0.0028; KO-SAL: 2.4 ± 0.43; KO-OXT: 1.70 ± 0.52, Kruskal-Wallis; KO-OXT vs WT-OXT: p>0.99, Dunn’s post-hoc) (**Fig. 4B**). The reduction in AGS was accompanied by a greater latency to seizure onset in *Fmr1*-KOs receiving OXT (Log-rank (Mantel-Cox) test = 6.07; p=0.01) (**Fig. 4C**). To test if acute OXT treatment has enduring effects comparable to those observed following early-life treatment, the same cohort of *Fmr1*-KO mice given acute OXT were reassessed for AGS 24 h and 15 d later. At the 24h time point, seizure behavior remained low in OXT-treated mice (30-60 min vs 24hr: p>0.99, Dunn’s post-hoc), but by 15 d post-treatment the AGS response returned to the robust expression (FM= 14.86, *p*= 0.0002, Friedman; KO-OXT (30-60 min): 0.20 ± 0.13; KO-OXT (1d): 0.50 ± 0.17; KO-OXT (15d): 2.60 ± 0.52; 30-60 min vs 15 d:**p=0.005, Dunn’s post-hoc) (**Fig. 4D**). These results demonstrated that in *Fmr1*-KOs, the anti-epileptic effects of adult OXT treatment are transient in nature, and provide further support for there being a critical developmental window for realizing an enduring reduction in AGS susceptibility with OXT treatment.

**Figure 4.**
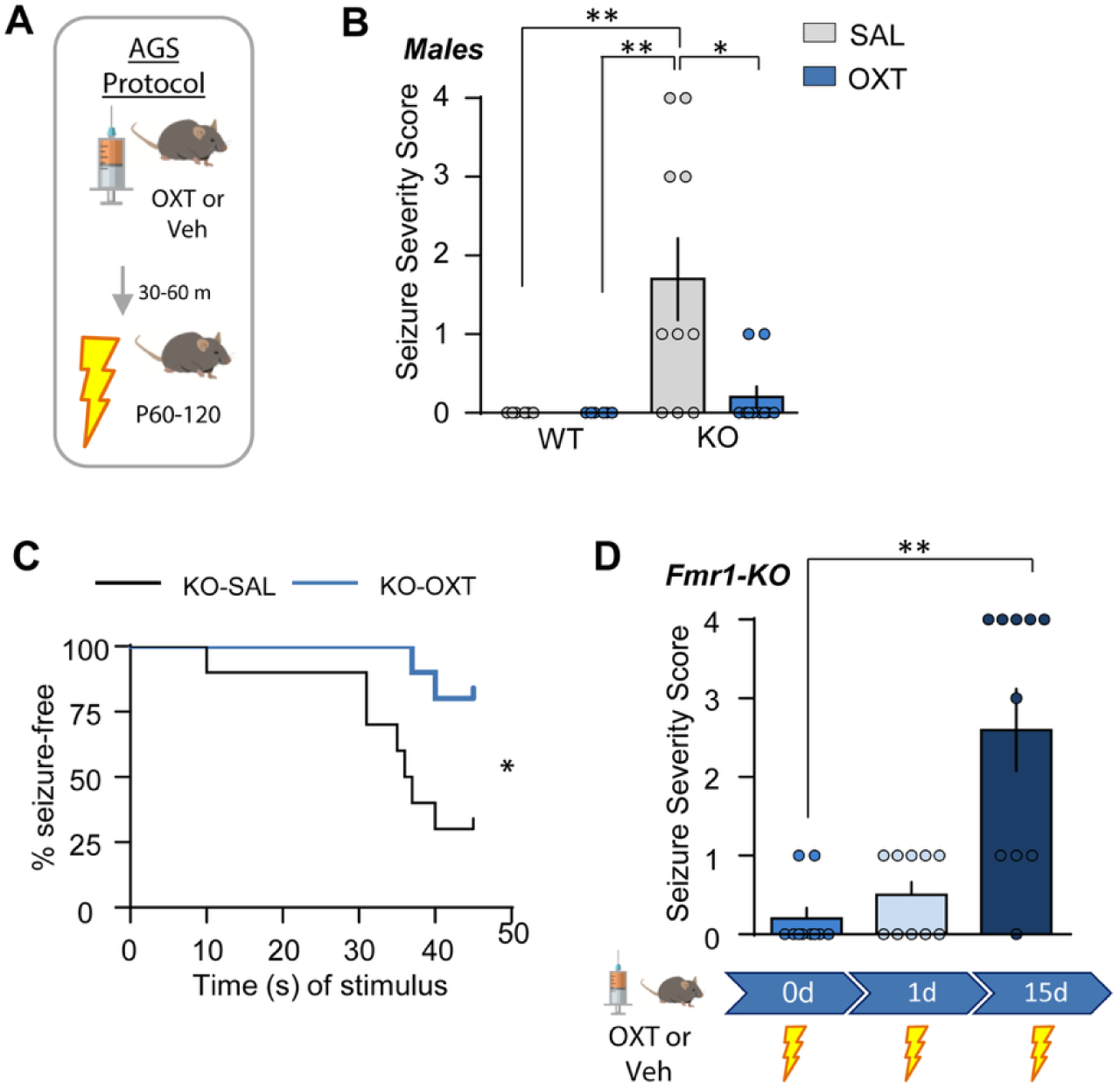
Acute oxytocin treatment produces a temporary attenuation of AGS in male *Fmr1*-KOs. (**A**) Schematic of AGS paradigm: adult mice were given an acute IP injection of oxytocin (OXT) or saline (SAL) and AGS testing began 30-60 min thereafter. (**B**) Wild type (WT) mice did not exhibit seizures independent of treatment. *Fmr1*-KOs given SAL displayed a broad range of seizure responses; seizure activity was markedly reduced in the OXT group (H=14.11, p=0.0028; WT-SAL n= 6, WT-OXT n=6, KO-SAL n=10, KO-OXT n=10). (**C**) The latency to seizure response was markedly lower in *Fmr1*-KOs given SAL than OXT (Log-rank= 6.07, p=0.01; Hazard ratio (KO-SAL)= 5.46 (95% CI= 1.50 to 21.8; KO-SAL n=7, KO-OXT n=2); note, the low “n” for the KO-OXT group was due to the majority of animals not seizing (see panel B). (**D**) Retesting of *Fmr1*-KOs given acute OXT demonstrated a resumption of robust seizure responses after 15 days (Friedman (RM)=14.86, p=0.0002; n/group =10). *Statistics*: Kruskal–Wallis (H) with Dunn’s post-hoc comparisons *(B)* and Log-rank (Mantel-Cox) test with hazard ratio (logrank) (*C)* indicated as: *p<0.05 and **p<0.01.

## DISCUSSION

Here we present novel evidence that OXT administration in the early postnatal period (P7-P13), but not during adolescence (P30-P36), significantly reduced the incidence and severity of AGS, and the latency to seizure onset, in male *Fmr1*-KOs tested in adulthood (>P60). Female KOs exhibited less severe seizure responses that are not affected by OXT administration. Furthermore, although a single acute OXT treatment reduced seizure responses in adult male KOs tested at 30 min and 1 day later, these effects were not evident 15 days later. Thus, adult OXT treatment did not have the long-lasting effects realized with OXT treatment as in the postnatal period. These findings highlight the potential of using early-life OXT treatment as an effective therapeutic intervention to eliminate the predisposition to seizures in male *Fmr1*-KOs and potentially in FXS. These results provide novel evidence for sex differences in both the expression of seizures and efficacy of OXT treatment in FXS model mice.

The maturation of neuronal circuits during postnatal brain development relies on a finely regulated balance of excitation and inhibition, and disruptions to this process have been implicated in neurodevelopmental disorders such as autism and FXS (Rubenstein and Merzenich, 2003; Belmonte and Bourgeron, 2006; Akerman and Cline, 2007; Paluszkiewicz et al., 2011). Studies have highlighted OXT’s involvement in facilitating the critical transition of GABAergic transmission to a progressively more hyperpolarized reversal potential during development, an event that is essential for establishing inhibitory control of neuronal excitability (Tyzio et al., 2006; Leonzino et al., 2016; Mahadevan and Woodin, 2016). This developmental change depends on the timely expression and functional upregulation of the main chloride exporter, KCC2, which lowers intracellular chloride levels and enables GABA_A_ receptor-mediated inhibition (Ben-Ari et al., 2007; Kaila et al., 2014; Mahadevan and Woodin, 2016; Virtanen et al., 2021). However, in *Fmr1*-KOs this shift is delayed, at least in the neocortex, such that the chloride reversal potential stays relatively depolarized on P10, but eventually normalizes by P14 (He et al., 2014; Tyzio et al., 2014). Studies in rodent autism models demonstrated that OXT receptor (OXTR) activation can modulate this shift by activating GPCR signaling to promote KCC2 expression and phosphorylation during postnatal development; the efficacy of OXT treatment to elicit this change aligns with our iOXT treatment timeline (Tyzio et al., 2006; He et al., 2014; Tyzio et al., 2014; Leonzino et al., 2016; Mahadevan and Woodin, 2016). Thus, it is possible that in the KOs, OXT treatment engages signaling that accelerates maturational changes in KCC2 during a critical developmental window, thereby normalizing network excitability. OXT administration outside this early postnatal period such as in adolescence -- when GABAergic signaling has already transitioned to its mature inhibitory state (Kilb, 2012) -- may have limited influence on circuit function, potentially explaining the lack of adolescent OXT treatment effects on AGS in adulthood. A goal for future work will be to directly test if, in the *Fmr1*-KOs, iOXT during the second postnatal week normalizes KCC2 expression and activity.

Although OXT may influence the polarity of GABA receptor currents during development, these processes would not be pertinent to the acute effects of OXT on AGS in adults. AGS progression relies upon hyperexcitability in several brain regions including auditory cortex, inferior colliculus and brainstem (Garcia-Cairasco et al., 1993; Chen and Toth, 2001; Fedotova et al., 2021). The G-protein coupled OXTR is broadly expressed including in forebrain and brainstem regions (Newmaster et al., 2020; Wang et al., 2025), and is localized to both excitatory and inhibitory synapses (Lin and Hsu, 2018). Although a range of OXTR functions have been described, largely in association with different G-proteins coupled to the receptor, there is good evidence that for some brain regions the predominant effect of OXT is direct activation of interneurons and, consequently, indirect negative modulation of excitatory cells (Lin and Hsu, 2018). This OXT action has been associated with changes in oscillatory network activity and reductions in the epileptogenic effects of pentylenetetrazol treatment (Erfanparast et al., 2017). Therefore, it is possible that acute increases in OXTR signaling prevent seizure progression by increasing inhibition within distributed neuronal networks. This could explain the transient effects of OXT treatment on AGS but raises questions about the duration of these acute-treatment effects.

Another intriguing aspect of the present results is the clear evidence for sex differences in both the predisposition for seizures and responsiveness to early OXT treatment. Preclinical FXS research has been strongly biased towards male subjects due to the X-linked inheritance of the disorder (Bagni et al., 2012). In humans, males with FXS typically have a full mutation on their one X chromosome, and thus lack *Fmr1* gene expression, whereas females typically have one mutated and one intact X chromosome with the latter mediating *Fmr1* expression (Hagerman and Hagerman, 2002). Overall, this leads to a reduction, but not elimination of FMRP expression, and a less severe cognitive and behavioral phenotype in females as compared to males carrying the mutation (Tsiouris and Brown, 2004; Huddleston et al., 2014; Bartholomay et al., 2019). However, in contrast to the human condition, the female mice used here are homozygous for the *Fmr1* KO and thus fully lack FMRP expression. It was therefore surprising that the AGS response in female KOs was markedly lower than that in males. Although prior studies have reported prominent seizure responses to AGS in both homozygous and heterozygous female *Fmr1*-KO mice (Yan et al., 2005; Musumeci et al., 2007; Ding et al., 2014; Guo et al., 2016), we observed only modest seizures and no detectable effect of OXT treatment. These findings are consistent with prior clinical studies which have shown that seizure susceptibility is inherently lower in females with FXS (Berry-Kravis et al., 2010).

Sex differences in seizure susceptibility could be a result of NMDAR operations differing between the sexes. Previous studies revealed that males rely on ion flux independent (i.e., metabotropic) NMDAR operations to trigger cytoskeletal changes that stabilize enduring synaptic plasticity in hippocampal field CA1 whereas females instead rely upon synaptic estrogen receptor-**α** (ER-α) for the same processes (Le et al., 2024). Similarly, in nucleus accumbens males rely on NMDARs for LTP of hippocampal afferents whereas females, instead, depend upon ER-α and voltage dependent calcium channels for the same functions (Copenhaver and LeGates, 2024). It has been suggested that aberrant NMDAR activity may play a role in the expression of seizures (Yen et al., 2004; Ghasemi and Schachter, 2011; Chen et al., 2021). Therefore, it is possible that sex differences in NMDAR involvement could contribute to the lower predisposition for seizures, including AGS, in females.

The male-only effects of OXT treatment observed in the present study also suggest that there are sex differences in the OXTRs or their neuronal functions. Studies have shown that OXTR expression is comparable in males and females for most brain regions, but more OXTR+ cells have are localized to female piriform cortex and hippocampal region CA2 cortex (Mitre et al., 2017; Mitre et al., 2018). Whether there are sex differences in receptor signaling per se is not known. However, it is noteworthy that the effects of applied OXT on synaptic transmission reportedly differ between neuronal groups and brain areas (Bakos et al., 2018) and, in individual studies, the nature of the response has been ascribed to the specific G-proteins coupled to the OXTR (Jurek and Neumann, 2018; Borroto-Escuela et al., 2022). This suggests that there may be sex differences in G-proteins associated with OXTRs in the network that propagates AGS such that the agonist (OXT) does not have the same effects on excitatory/inhibitory balance in females as it does in males. In support of this idea, it was recently reported that the OXTR agonist Thr4,Gly7-oxytocin (TGOT) reduces intrinsic neuronal excitability in juvenile male rats but has no effect in age-matched females (Rajan Narattil et al., 2026).

Together, the present findings provide novel evidence that early-life OXT treatment robustly attenuates seizure responses in adult male *Fmr1*-KO mice and that females, which exhibit milder seizure responses, are not responsive to the same treatments. In males, the enduring antiepileptic effects of OXT treatments are developmentally restricted, as neither delayed nor acute OXT administration produces long-lasting anticonvulsant effects. These results highlight the importance of early postnatal intervention and identify treatment options for FXS individuals vulnerable to seizures.

## FUNDING AND ACKNOWLEDGEMENTS

This work was supported by NIH NICHD Grants HD101642 and HD101642-S1, and NINDS training grant T32 NS04554.

